# Odorant representations indicate nonlinear processing across the olfactory system

**DOI:** 10.1101/2022.04.15.488302

**Authors:** Jesús Olivares, Patricio Orio, Viktor Sadílek, Oliver Schmachtenberg, Andrés Canales-Johnson

## Abstract

The olfactory system comprises intricate networks of interconnected brain regions that process information across both local and long-range circuits to extract odorant identity. Similar to pattern recognition in other sensory domains, such as the visual system, recognizing odorant identity likely depends on highly nonlinear interactions between these recurrently connected nodes. In this study, we investigate whether odorant identity can be distinguished through nonlinear interactions in the local field potentials (LFPs) of the olfactory bulb and telencephalic regions (Vv and Dp) in anesthetized rainbow trout. Our results show that odorant identity modulates complex information-theoretic metrics, specifically information sharing and redundancy, across these brain areas, indicating nonlinear processing. In contrast, traditional linear connectivity measures, such as coherence and phase synchrony, showed little or no significant modulation by odorants. These findings suggest that nonlinear interactions encoded by olfactory oscillations carry crucial odor information across the teleost olfactory system, offering insights into the broader role of nonlinear dynamics in sensory processing.

## INTRODUCTION

Sensory systems, particularly the olfactory system, are composed of intricate networks of interconnected brain regions that process information across both local and long-range circuits. Olfaction is notable for its reliance on distributed processing, with the olfactory bulb transmitting sensory input to higher-order brain regions where odor information is integrated and interpreted (Mori et al., 1999; Haberly, 2001). However, due to the complexity of this system and the particular nature of olfactory objects, the precise mechanisms by which different types of neural dynamics encode odorant information across brain regions remain poorly understood (Fulton et al., 2024; Rokni and Ben-Shaul, 2024). In olfactory processing, much of the early work has focused on oscillatory activity within local regions, with specific rhythms such as theta (4-8 Hz), beta (12-30 Hz), and gamma (30-100 Hz) implicated in various aspects of sensory encoding (Gelperin, 2006; Kay et al., 2009; van Wingerden et al., 2010; Martin and Ravel, 2014). Oscillations are thought to facilitate communication between neurons and between brain areas by synchronizing neural activity over time, a process that could theoretically enhance the transmission of sensory information. However, the degree to which these oscillatory interactions can encode and differentiate between different odorants remains uncertain(Kay et al., 2009). This raises a fundamental question: Is the functional connectivity between olfactory regions modulated by the identity of odorants, and if so, what type of neural dynamics plays a key role in this modulation?

The traditional view has been that oscillatory synchronization, reflected in measures like coherence and phase synchrony, supports communication across brain areas by temporally aligning neural activity (Singer, 1999, 2018). These linear measures focus on the periodic nature of brain activity, assessing how closely signals in different regions oscillate in unison. Oscillatory coherence has been observed in a wide range of sensory modalities, including vision and audition, and similar phenomena have been described early on in the olfactory system (Adrian, 1942; Bressler and Freeman, 1980). It is now well established that odorant exposure can modulate the power and coherence of oscillatory activity in the olfactory bulb and higher olfactory brain regions in mammals and analogous structures in invertebrates(Laurent, 2002; Martin and Ravel, 2014). Yet, while these oscillations reflect some aspects of sensory processing, their precise role in long-range communication between olfactory regions and their ability to encode the complex identity of distinct odorants remain unresolved.

Recent advances in neuroscience have highlighted the limitations of focusing solely on measures that quantify interactions within the same frequency band, such as coherence and Granger causality, as these measures can be strongly influenced by linear information transmission (Srinath and Ray, 2014; Vinck et al., 2023; Pesaran et al., 2018; Vinck et al., 2025; Schneider et al., 2021). Such linear information transmission can emerge from anatomical projections in a sending to a receiving area (Schneider et al., 2021). However, interareal communication may be better described by nonlinear dynamics, which capture more complex, often arrhythmic interactions across regions(Baird, 1986). These dynamics are best quantified by modern information-theoretic metrics such as information sharing and information redundancy. These nonlinear measures focus on how information is distributed and shared between areas, regardless of whether that information is carried by rhythmic signals (Vinck et al., 2023). The concept of information sharing refers to the amount of mutual information between two or more brain regions. When brain areas share high levels of information, it suggests that these regions are working together to process the same aspects of sensory input, such as an odorant’s identity. Similarly, information redundancy captures whether the encoded trial-by-trial information about the stimuli is indeed the *same* in different brain areas, providing insight into the robustness of neural communication (Roberts et al., 2025). These nonlinear metrics offer a more nuanced view of brain functional connectivity, capturing interactions that linear measures such as coherence and phase synchrony, which quantify interactions between areas only at the same frequency, might miss (Imperatori et al., 2019; Canales-Johnson et al., 2020c, 2023; Vinck et al., 2023; Gelens et al., 2024; Vinck et al., 2025).

This study addresses a central question in olfactory neuroscience: What type of dynamic interaction between brain areas best encodes odorant identities? Specifically, we compare linear functional connectivity measures (such as coherence and phase synchrony) with nonlinear measures (information sharing and redundancy) to determine which type of functional connectivity measure is modulated by the identity of different odorants. By investigating both linear and nonlinear dynamics, we aim to provide a more comprehensive understanding of how the brain encodes olfactory information across long-range networks. To explore this, we used an olfactory model, anesthetized juvenile rainbow trout, in which the olfactory organ is not connected to the respiratory system, to isolate neural activity related to odorant processing from potential confounds such as respiration, learning, and behavior(Olivares and Schmachtenberg, 2019).This model offers a unique advantage: It allows us to study the neural dynamics of olfaction without the interference of sniffing, a behavior that can complicate the interpretation of oscillatory signals in mammals. Using local field potential (LFP) recordings from the olfactory bulb and the telencephalic regions Vv and Dp together with the electroolfactogram, we compared how different odorants modulate both linear and nonlinear interactions between these regions.

## MATERIALS AND METHODS

### Animals, anesthesia, and surgery

Juvenile rainbow trout, *Oncorhynchus mykiss*, were obtained from a local hatchery (Piscicultura Río Blanco, Los Andes, Chile), and kept at 16°C in aquariums with dechlorinated and filtered water in the animal facility of the Faculty of Sciences of the Universidad de Valparaíso, for up to six months, with a light / dark regime of 12:12 hours. The animals were fed daily with fish meal pellets. For the experiments, specimens of undetermined sex and a total length of 20 ± 2 cm were selected. The procedures applied in this study were approved by the Bioethics Committee of the Universidad de Valparaíso (Code: BEA 100-2017) and are in accordance with the guidelines of the National Research and Development Agency (ANID) of Chile. Before the experiments, the fish were anesthetized with 10 mg/l MS-222 (ethyl 3-aminobenzoate methanesulfonate) and immobilized with an intramuscular injection of gallamine triethiodide (Flaxedil; 0.03 mg / 100 g body weight), until balance and responsiveness were lost. Immobilized fish were wrapped in moistened tissue and placed in a container of synthetic sponge. The gills were irrigated through an oral tube with a constant flow of cooled aerated water (10°C) containing 10 mg/l of MS-222 (pH 7.4). Before surgery, all dissection material was impregnated with lidocaine hydrochloride (2%), which was also applied directly on the surface of the animal’s skin. To gain access to the fish’s olfactory organ, the skin and connective tissue covering the entrance to the nostril was removed. To access the brain, the cranial vault was opened under a stereomicroscope with a transverse section through the midline of the head at eye level. Then, two cuts were made perpendicular to the first, and the skull was carefully opened until the brain was exposed. From this point on, the brain was periodically irrigated with artificial cerebrospinal fluid (in mM: 124 NaCl, 2.69 KCl, 1.25 KH_2_PO_4_, 2.0 MgSO_4_, 26 NaHCO_3_, 2.0 CaCl_2_, 10 glucose, pH 7.4). The vitality of the fish was monitored by checking the blood flow in the vessels of the *tela coroidea* covering the telencephalon.

### Odor preparation and stimulation

The water to which the trout were adapted in the animal facility (TW, trout water) was used as negative odor control, and to dilute the odorant mixtures. Four different types of odor stimuli were applied in the study: (1) A synthetic mixture of five L-amino acids (AA): Serine, cysteine, aspartate, lysine, and alanine; a mixture of four synthetic bile salts (BS): Deoxycholic acid, sodium taurocholate, lithocholic acid and taurolithocholic acid; beta-phenylethyl alcohol (PEA) as single odorant stimulus, and an extract of trout skin (SE). Amino acids represent food odors to fish; bile salts are considered to be social odors and PEA is a compound that is used as a fishing lure and as synthetic odorant in zebrafish studies (Nikonov and Caprio, 2004; Rolen and Caprio, 2008; Calfún et al., 2016). Finally, the extract of trout skin is a natural odorant mixture, likely composed of hundreds of compounds, containing alarm pheromones among others (Nikonov and Caprio, 2004; Wisenden et al., 2009; Mathuru et al., 2012; Valdés et al., 2015).

The amino acids and bile salts were prepared as stock solutions at a concentration of 100 mM and subsequently diluted in TW to the final concentration. Skin extract was prepared from the macerated skin of five juvenile trout in 100 ml of TW, filtered, and aliquoted as stock solution. The stock was frozen at -80°C and used throughout the study. All chemicals were purchased from Sigma-Aldrich (Santiago, Chile). Approximately equipotent odor stimuli were defined by diluting BS and SE stocks to concentrations generating the same electroolfactogram amplitude as 100 µM of AA, yielding 700 µM for the BS mixture and a dilution of 1: 500 for the SE stock. For PEA, equipotent values could not be obtained, since responses remained smaller up to concentrations above 1 mM. The olfactory stimuli were administered automatically using a custom-built computer-controlled picospritzer connected to the gravity-fed open perfusion system constantly irrigating the olfactory organ.

### Electrophysiological recordings

We recorded the electroolfactogram (EOG) in parallel with the local field potentials (LFP) in the olfactory bulb (OB) and two telencephalic regions, the ventral nucleus of the ventral telencephalon (Vv) and the dorsal posterior zone of the telencephalon (Dp) (**Figure 1A**). All signals were acquired with WinWCP software version 4.3.8 (John Dempster, University of Strathclyde, UK) and analyzed using Clampfit 10.3. (Molecular Devices). The EOG was recorded from the olfactory rosette with an Ag/AgCl electrode located directly above the central raphe. An identical reference electrode was placed on the skin of the head, and a ground electrode connected to a fin. The EOG signals were amplified with a differential AC amplifier (Warner DP-301, Warner Instruments, USA) and digitized with a PCI-6221 A/D interface (National Instruments, USA) at a sampling rate of 1 kHz. For each recorded brain sector, two tungsten electrodes were used (FHC tungsten microelectrode, tip diameter 3 µm, impedance 11-13 MΩ), one at the recording site and the other as a reference, positioned in the cerebrospinal fluid of the exposed cranial cavity. In the olfactory bulb, the electrodes were positioned within the mitral cell layer, about 150 µm deep with respect to the dorsal surface of the bulb. Given the odotopy of the olfactory bulb, the position of the recording electrode was adjusted depending on the stimulus type, based on an odotopy map reported in (Bazáes et al., 2013). The electrode used to record Vv was positioned in the anterior third of the animal’s telencephalon at approximately 500 µm from OB and a depth of 150 µm, entering through the central sulcus between the hemispheres. The electrode recording the telencephalic region Dp was positioned in the posterior third of the telencephalon, at 500 µm distance from the optic tectum (OT) and a depth of about 450 µm. All recordings were made in ipsilateral brain structures with respect to the stimulated olfactory organ. Each recording protocol had a duration of 50 seconds, which included the application of the olfactory stimulus or control after the first 10 second with a duration of 1 s. Twenty consecutive trials were performed for each stimulus type. The LFP electrodes were connected to the headstages of an AC amplifier (AM Systems Model 1800), band-pass filtered between 1 and 100 Hz and digitized at 1 kHz.

**Figure 1.**
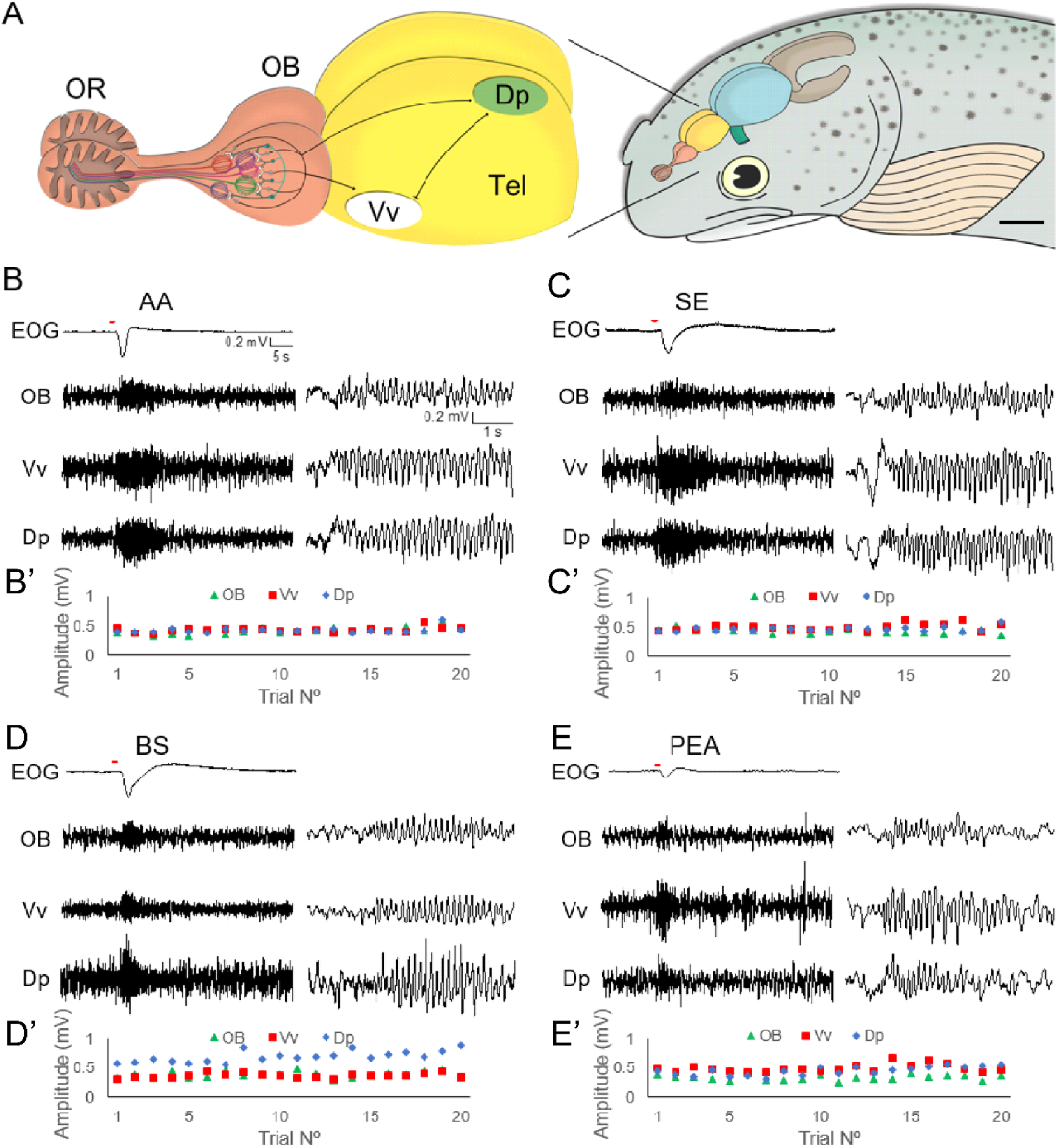
Recording sites and raw simultaneous recordings of the EOG and LFP oscillations from the olfactory bulb (OB), the ventral nucleus of the ventral telencephalon (Vv) and the dorsal posterior zone of the telencephalon (Dp). (**A**) Schematic drawing of the rainbow trout forebrain. Olfactory information and LFP oscillations of different frequency bands are transmitted in parallel from the olfactory bulb to higher olfactory areas of the telencephalon, notably Vv and Dp. Scale bar: 3 mm. (**B-E**) Responses to an amino acid (AA, 100 μM) and a bile salt (BS, 700 μM) mix, conspecific skin extract (SE, diluted 1:500) and beta-phenylethyl alcohol (PEA, 100 μM) were recorded simultaneously from OB, Vv and Dp, showing prominent and ostensibly synchronous oscillatory activity during odor stimulation. The sets of traces to the right display details on a shorter timescale. (**B’-E’**) Maximum amplitude of oscillation reached in each of the 20 applications of odorant. Oscillatory response amplitudes remained stable for 20 trial repetitions in each olfactory brain area, showing no sign of sensory adaptation.

### Signal processing and frequency analysis

Signals were low pass filtered at 50 Hz (3 pole Butterworth filter) and downsampled to 200 Hz. For each experiment, a frequency spectrum was calculated from the 10 s following each of the stimulus exposures, and the spectra were averaged. Similarly, baseline spectra were calculated for the 10 s prior to stimulus application and averaged. The average baseline spectra were subtracted from the post-stimulus. Spectrograms were calculated using a continuous wavelet transform (CWT) with the complex Morlet wavelet (σ=5). The CWT is defined as:

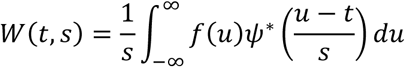

Where *s* is the scale parameter, *t* the position parameter, *f*() the signal function, *ψ*() the wavelet function and the asterisk represent the complex conjugate. The complex Morlet wavelet is defined as:

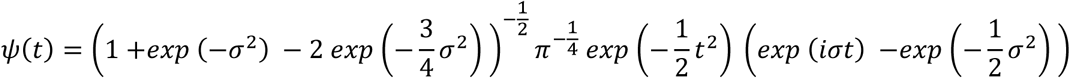

With central frequency ∼*σ*. The scaleogram, analog to the spectrogram, is defined as the square of the amplitude of the CWT:

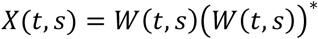

The scale is related to the period (T) and frequency (*f*) in the following relationship:

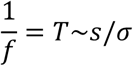

The spectrogram amplitudes were expressed as the z-score by subtracting, at each frequency, the mean of the amplitudes of the baseline spectrogram and dividing by its standard deviation.

### Coherence analysis

Coherence *C*_*XY*_ between signals *X* and *Y* was calculated as:

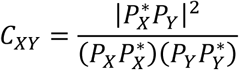

being *P*_*X*_ and *P*_*Y*_ the spectral power density estimators of signals *X* and *Y*, respectively. The asterisk denotes the complex conjugate. The estimation of spectral power was carried out with Welch’s method, using windows of 500 samples and an overlap of 350 samples. Like the frequency spectra, the coherence of the 10 s prior to the stimuli was subtracted from the coherence calculated in the 10 s after the stimuli. Spectral and coherence calculations were performed in Python language with the routines provided by the Scipy package. Wavelet spectrograms were performed using Python code freely available at https://github.com/patoorio/wavelets.

### Spectral decoding

In addition to the inferential statistical approach, spectral decoding was applied to the time-frequency data. This was done by performing Multivariate Pattern Analysis (MVPA) and univariate pattern analysis. MVPA decoding holds an advantage over univariate decoding as it offers more spatially sensitive dependent measures, demonstrating that information is present in activity patterns across brain regions (Fahrenfort et al., 2018a; Hebart and Baker, 2018). The ADAM toolbox (Fahrenfort et al., 2018b) was used on raw LFP data, which was transformed to time-frequency charts with similar settings epochs: 0 to 10 seconds after stimuli presentation, 1–40 Hz. For the MVPA decoding, we used as a feature the single-trial spectral power values per frequency bin and time points across the three olfactory regions (OB, Vv, Dp) **(Figure 3B)**. For the univariate decoding, we used as a feature the single-trial spectral power values per frequency bin and time point, performed in each region separately **(Figure 3C)**. For both analyses, we performed multiclass decoding, where the four classes of stimuli were separated simultaneously.

It is essential to keep a balanced number of trials between conditions when performing a multivariate decoding analysis since design imbalances may have unintended effects (Fahrenfort et al., 2018b) on the linear discriminant analysis (LDA; the classification algorithm used here) and area under the curve accuracy metric (AUC; the accuracy performance metric used here). To keep a balanced number of trials across conditions, we randomly selected and discarded trials when necessary (undersampling). We quantified classifiers’ accuracy performance by measuring the AUC of the receiver operating characteristic (ROC), a measure derived from signal detection theory that is insensitive to classifier bias. AUC corresponds to the total area covered when plotting the cumulative true positive rates against the cumulative false positive rates for a given classification task. Thus, finding above-chance performance indicates that there was information contained in the neural data that the classifier decoded based on the stimulus features of interest.

LFP epochs time-locked to odorant presentation were classified according to their type (i.e., SE, AA, BS). Next, a backward decoding algorithm using the odorant category was applied according to a standard 10-fold cross-validation scheme. Thus, we separated data sets for training and testing in each trout; the classifier was trained on 90% of trials and tested on the remaining 10%, repeating this procedure time and averaging across folds to obtain an AUC score. An LDA was used to discriminate between all odorant classes (multiclass decoding) after which classification accuracy was computed as the AUC, a measure derived from signal detection theory. AUC scores were tested per time point with double-sided t-tests across participants against a 50% chance level. These t-tests were double-sided and corrected for multiple comparisons using cluster-based 1000-iteration permutation tests (Maris and Oostenveld, 2007) with a standard cut-off p-value of 0.05. This procedure yields time clusters of significant above-chance classifier accuracy, indicative of information processing.

### Measurement of phase synchronization

We quantified phase locking between pairs of electrodes to measure dynamical interactions among electrode signals oscillating in the same frequency range. Phase synchronization analysis proceeds in two steps: the estimation of the instantaneous phases, and the quantification of the phase locking. To obtain the instantaneous phases, *φ*, of the neural signals, we used the Hilbert transform. The analytic signal *ξ*(*t*) of the univariate measure *x*(*t*) is a complex function of continuous time defined as:

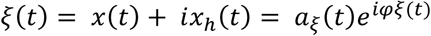

where the function is the Hilbert transform of:

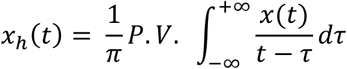

P.V. indicates that the integral is taken in the sense of Cauchy principal value. Sequences of digitized values give a trajectory of the tip of a vector rotating counterclockwise in the complex plane with elapsed time. The vector norm at each digitizing step *t* is the instantaneous amplitude

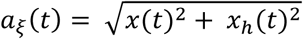

Similarly, the complex argument of analytic signal is the instantaneous phase *φ*_*x*_(*t*).

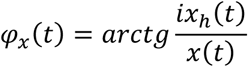

The instantaneous phase, although defined uniquely for any signal to which the Hilbert transform can be applied, is challenging to interpret for broadband signals. For this reason, a standard procedure is to consider only narrow-band phase synchronization by estimating an instantaneous phase for successive frequency bands, which are defined by band-pass filtering the time series(Le Van Quyen et al., 2001). Thus, we band-passed filtered the LFP signals in multiple consecutive 1 Hz-wide frequency bins from 6 to 10 Hz using a zero-phase shift non-causal finite impulse filter.

### Phase locking quantification: weighted phase lag index (wPLI)

Phase synchronization can be considered as an LFP/EEG measure of oscillatory coupling between neuronal populations(Fries, 2005; Imperatori et al., 2019). The Phase Lag Index (PLI)(Stam et al., 2007) attempts to minimize the impact of volume conduction and common sources inherent in EEG data by averaging the signs of phase differences, thereby ignoring average phase differences of 0 or 180 degrees. This is based on the rationale that such phase differences are likely to be generated by volume conduction of single dipolar sources. Despite being insensitive to volume conduction, PLI has two important limitations: first, there is a strong discontinuity in the measure, which causes it to be maximally sensitive to noise; second, when calculated on small samples, PLI is biased toward strong coherences (i.e., it has a positive sample-size bias). Formally, the PLI is defined as the absolute value of the sum of the signs of the imaginary part of the complex cross-spectral density *S_xy_*of two real-valued signals *x*(*t*) and *y*(*t*) at time point or trial *t*:

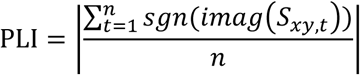

The Weighted PLI measure (wPLI) (Vinck et al., 2011) addresses the former problem by weighting the signs of the imaginary components by their absolute magnitudes:

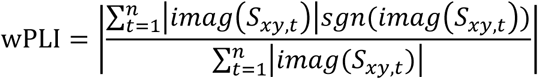

The PLI sample size problem can be addressed by considering the same number of trials per condition in the phase coherence analysis. Further, wPLI represents a dimensionless measure of connectivity that is not directly influenced by differences in spectral or cross-spectral power. The wPLI index ranges from 0 to 1, with a value of 1 indicating perfect synchronization (phase difference is perfectly constant throughout the trials) and value 0 representing the total absence of synchrony (phase differences are random). Temporal evolution of wPLI was calculated using a 500 ms sliding window with 2 ms time step, i.e. with a 96% overlap between two adjacent windows. wPLI charts are expressed in z-scores (standard deviation units) relative to the pre-stimulus period (1 s) which was regarded as a baseline.

### Information sharing between recording sites

We quantified the information sharing between electrodes by calculating the weighted symbolic mutual information (wSMI). This index estimates to which extent two LFP/EEG signals exhibit non-random joint (i.e., correlated) fluctuations. Thus, wSMI has been proposed as a measure of neural information sharing (King et al., 2013) and has three main advantages. First, it is a rapid and robust estimate of signals’ entropy (i.e., statistical uncertainty in signal patterns), as it reduces the signal’s length (i.e., dimensionality) by looking for qualitative or ‘symbolic’ patterns of increase or decrease in the signal. Second, it efficiently detects high nonlinear coupling (i.e., non-proportional relationships between neural signals) between LFP/EEG signals, as it has been shown with simulated (Imperatori et al., 2019) and experimental EEG/LFP data (Canales-Johnson et al., 2020a). Third, it rejects spurious correlations between signals that share a common source, thus prioritizing non-trivial pairs of symbols.

We calculated wSMI between each pair of electrodes and for each trial, after transforming the LFP signal into a sequence of discrete symbols defined by ordering of k time samples with a temporal separation between each pair (or τ). The symbolic transformation is determined by a fixed symbol size (k = 3, i.e., 3 samples represent a symbol) and the variable τ between samples (temporal distance between samples), thus determining the frequency range in which wSMI is estimated (King et al., 2013; Sitt et al., 2014). We chose τ = 32. The frequency specificity f of wSMI is related to k and τ as follows:

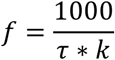

This formula, with a kernel size k of 3 and τ values of 32, produced a sensitivity to frequencies in the range ∼6 to 10 Hz in which the peak of oscillatory activity is observed in the Power Spectral Density (PSD) (**Figure 2**). wSMI was estimated for each pair of transformed LFP signals by calculating the joint probability of each pair of symbols. The joint probability matrix was multiplied by binary weights to reduce spurious correlations between signals. The weights were set to zero for pairs of identical symbols, as these could have been elicited by a unique common source, and for opposite symbols (i.e., of in opposite direction), as these could reflect the two sides of a single electric dipole. The following formula calculates wSMI:

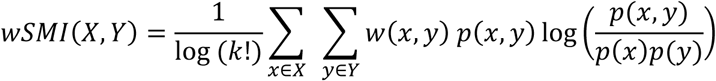

**Figure 2.**
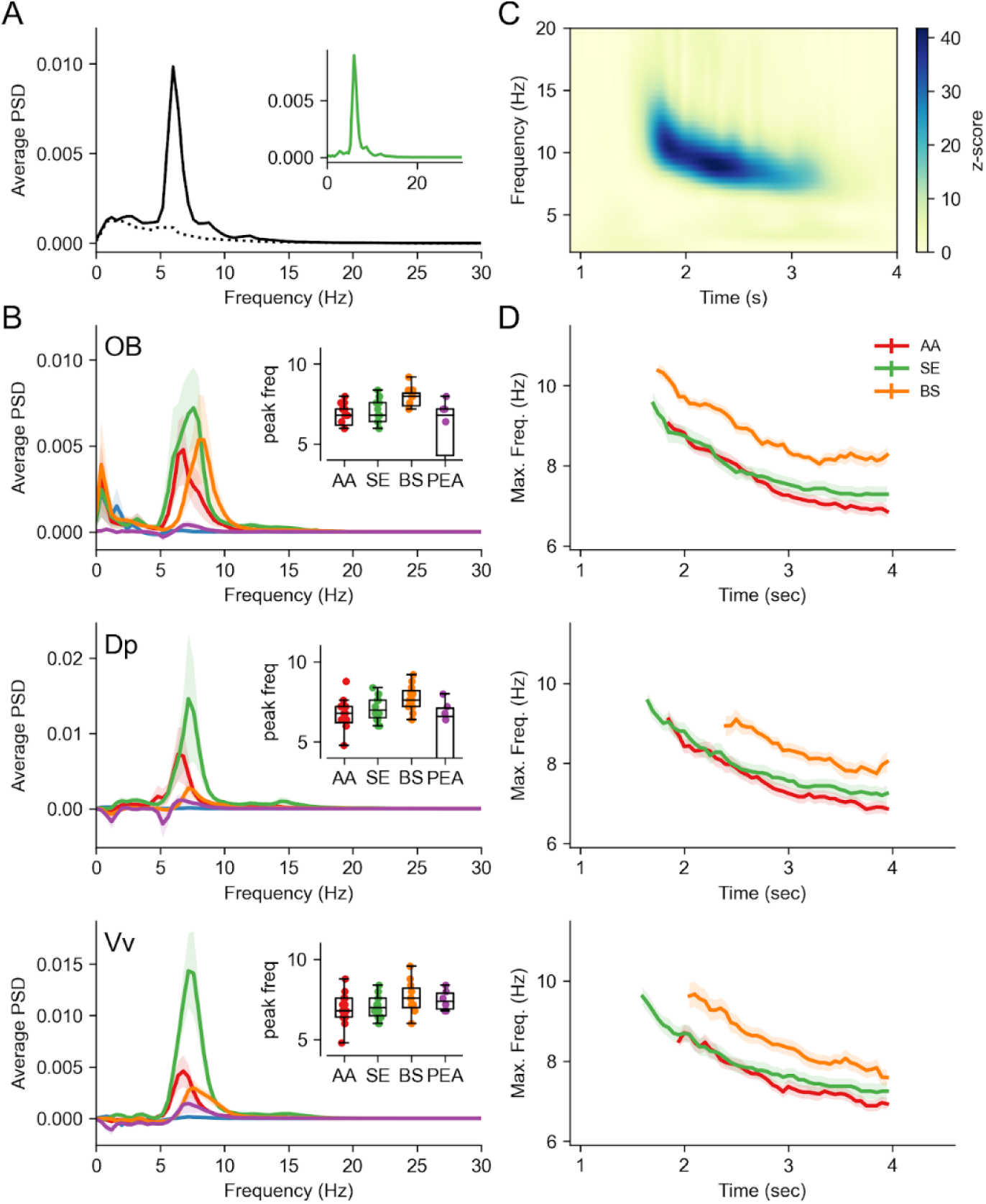
Oscillatory characteristics of neural response to odorants. (**A)** Average frequency spectrum of the LFP response recorded in Vv, upon exposure to the AA mix. The dotted line is the spectrum of the recording prior AA exposure (10 s) and the continuous line corresponds to the 10 s after exposure to AA. The spectra of 20 repetitions were averaged. The inset shows the net response to the odorants (baseline subtracted). (**B)** Average of baseline-subtracted spectra of the responses from OB, Vv, and Dp when stimulated with AA, trout skin extract (SE), bile salts (BS) and PEA, in addition to trout water control (TW; blue line). The average spectra of 11-14 experiments per condition are shown, with the mean as continuous line and the standard error as colored shade. The inset shows the peak of the response spectra. (**C)** Wavelet spectrogram of oscillations recorded from the OB upon exposure to SB at time t = 0 s. The data are expressed as z-score with respect to the baseline activity at each frequency during the 10 s prior to the stimulus. As in (A), it represents an average of 20 trials repeated in the same specimen with the same stimulus. (**D)** Average (lines) and standard error (shadow) of the maximum frequency at each time point, according to the wavelet spectrogram. The average was calculated only at times when more than half of the experiments showed a z-score higher than 5 in any frequency within the 5-20 Hz range, thus the different durations of the responses. Maxima were obtained from baseline-subtracted spectrograms.

**Figure 3.**
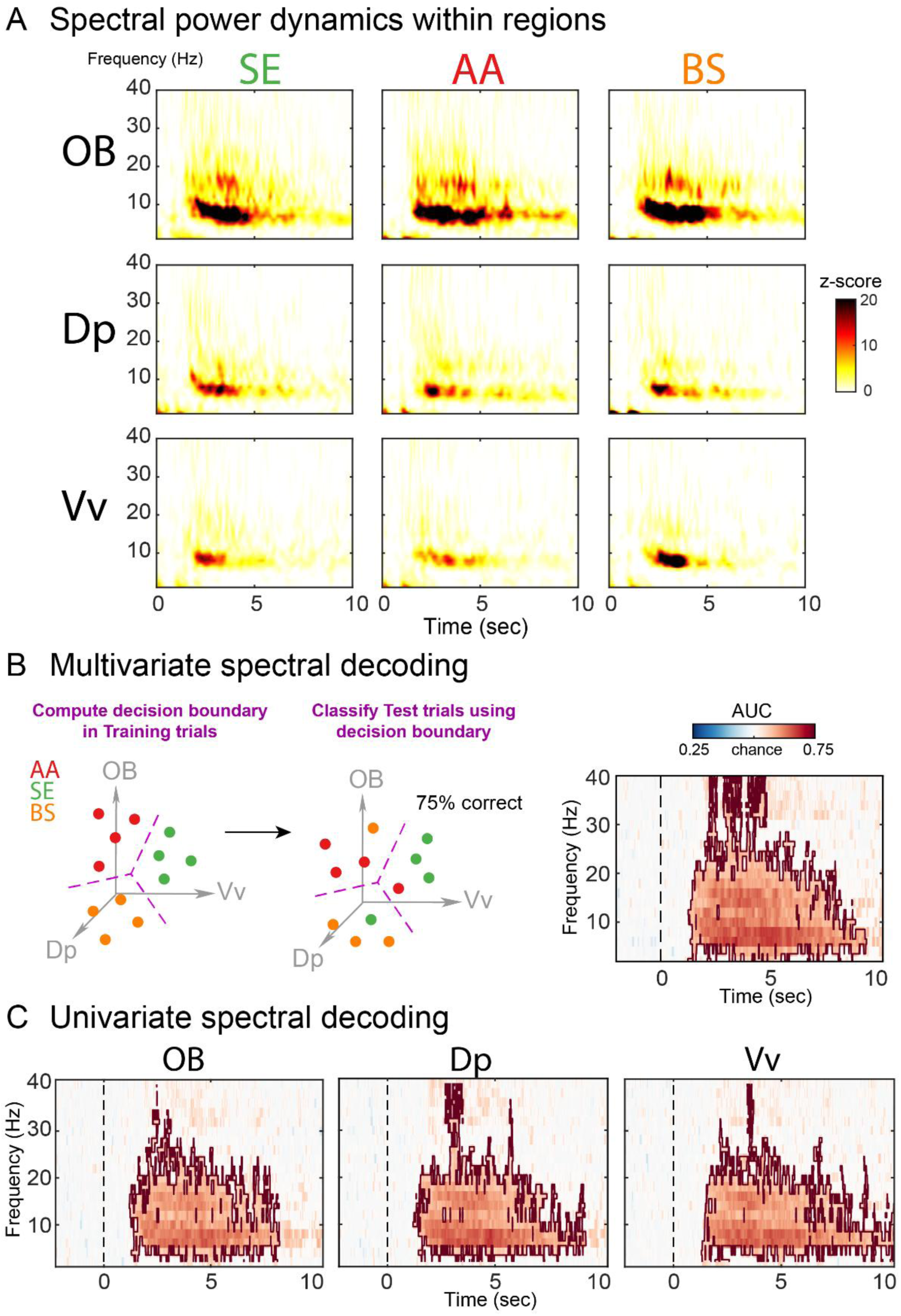
Spectral decoding in OB, Vv and Dp. **(A)** Temporal dynamics of spectral power. Time-frequency charts were averaged across individuals per odorant and region. **(B)** Multivariate spectral decoding. Each region is considered as a separate dimension, or “feature”, in a N-dimensional space, depicted by the 3-dimensional axes conformed by OB, Vv and Dp regions (in grey). Each trial-wise odorant presentation (AA in red, SE in green, and BS in orange) produces a pattern that occupies a point in a 3-dimensional neural activation space. A linear classifier (LDA) learns a way to transform this high-dimensional space into a new one in which the channel patterns associated with each odorant are separable by a decision boundary (left panel). LDA assigns an odorant label for the training data based on the position of the activity patterns relative to the decision boundary. The performance of the classifier is then a function of the accuracy of its label assignments (e.g., percentage correct, middle panel). This procedure is performed at each time point and frequency and a cluster-based permutation test (p<0.05; brown boundary) was performed to determine significant decoding above chance (see Materials and Methods). **(D)** Univariate spectral decoding. Same procedure as described above but for OB, Vv and Dp regions separately.

Here, x and y are symbols present in signals X and Y respectively; w(x,y) is the weight matrix and p(x,y) is the joint probability of co-occurrence of symbol x in signal X and symbol y in signal Y. Finally, p(x) and p(y) are the probabilities of those symbols in each signal and k! is the number of symbols used to normalize the mutual information by the signal’s maximal entropy. Temporal evolution of wSMI was calculated using a 500 ms sliding window with 2 ms time step, i.e., with a 96% overlap between two adjacent windows.

### Co-Information between recording sites

For each trial, after band-pass filtering the LFP signals between 6 to 10 Hz, we computed the time-resolved spectral power by computing the square of the instantaneous amplitude *a*_*ξ*_(*t*) using the Hilbert transform procedure described above. Using this resulting signal (i.e., the time-resolved spectral power), we then computed the co-Information (co-I) using the Gaussian Copula Mutual Information method (GCMI) (Ince et al., 2017). The co-information (co-I) has been calculated in the following way:

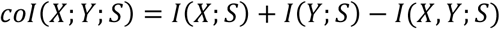

For each time point, *I(X;S)* corresponds to the mutual information (MI) between the signal at recording site *X* and stimuli class *S*. *I(Y;S)* corresponds to the MI between the signal at recording site *Y* and stimuli class *S*. Finally, *I(X,Y;S)* corresponds to the MI between stimuli class *S* combining signals from recording sites *X* and *Y*. This way, co-information was computed for each pair of odorants between all three pairs of recording sites. Positive co-information means that signals between recording sites contain redundant or overlapping information about the stimuli. Negative co-information corresponds to synergy between the two variables. This means that the mutual information when considering the two variables jointly is larger than when considering the variables separately.

### Statistical Analysis

Statistical analyses were performed using MATLAB (2016a), Jamovi (Version 0.8.1.6) [Computer Software] (Retrieved from https://www.jamovi.org) (open source), and JASP Team (2018; JASP; version 0.8.4 software) statistical software.

## RESULTS

### General description of the electrical responses

To retest the still unanswered hypothesis of whether odor information is carried by LFP oscillations from the olfactory bulb to higher olfactory centers, we recorded olfactory responses as reflected by variations of the LFP to four different types of odors: Two synthetic mixtures, amino acids, and bile salts, one natural mixture resulting from the mechanical maceration of conspecific skin, and the individual synthetic odorant PEA. Recordings were obtained simultaneously from the olfactory epithelium, the olfactory bulb, and two principal olfactory centers of the telencephalon: The subpallial ventral nucleus of the ventral region (Vv) and the pallial posterior dorsal region (Dp) (**Figure 1A**). Odor stimulation, as opposed to trout water (TW, control), always generated a dose-dependent negative EOG, and a transient and near-simultaneous increase in LFP oscillation amplitudes in the olfactory bulb and the telencephalic olfactory areas, lasting 10 to 15 s (**Figure 1B-E**). To test if repeated trials affected the LFP oscillations through sensory adaptation, sensitization, or learning-related mechanisms, we compared the responses of 20 consecutive trials, spaced 50 s apart, to the same stimuli (**Figure 1B’-E’**). Interestingly, the amplitudes of the oscillatory responses and the response envelopes remained unaltered throughout the trial sequence in the OB, and the telencephalic areas Vv and Dp. These findings allowed us to pool the data from several consecutive trials for subsequent analyses.

### Oscillatory power in the OB, and telencephalic areas Vv and Dp

We characterized the frequency-domain characteristics of the electrophysiological recordings and the responses to odorants, using the Welch periodogram method (**Figure 2A, B**). To improve the signal-to-noise ratio, we averaged the spectra across trials of the same experiment. Figure 2A shows the average spectrum calculated from a 10 s window before (dotted line) and after (continuous line) exposure to the amino acid mixture (AA), while recording in Vv. The baseline activity contains prominent low-frequency (<7 Hz) activity, and thus the baseline spectrum was subtracted from the odorant response spectrum (**Figure 2A, inset**). This subtraction was performed on all data shown in **Figures 2 and 3**. **Figure 2B** shows the average of the spectra for all experiments and their recordings in OB, Vv and Dp, while exposed to AA, SE, BS and PEA. Controls with trout water are shown in blue. AA, SE and BS evoked strong oscillatory responses in the recorded brain areas in the 5-9 Hz band, albeit with different magnitudes. The response to PEA, however, failed to generate robust oscillation in most experiments. We hypothesize that this is due to PEA, a component of floral odors, being a novel odorant for the trouts, with no or little biological meaning. Therefore, the responses to PEA were excluded from the following frequency-domain analyses.

The frequency of the maximum amplitude for each experiment is displayed in the insets. Notably, the response to BS appears to occur at a higher frequency on average than the response to AA and SE. However, only at OB and Dp a significant difference in the maximum frequencies was detected (Kruskal-Wallis test: OB H=6.43, p=0.04; Dp H=7.84, p=0.02; Vv H=1.75, p=0.42), and the difference was only found between AA and BS (Dunn’s posthoc test: OB AA/SE p=0.52, AA/BS p=0.013, SE/BS p=0.06; Dp AA/SE p=0.21, AA/BS p=0.005, SE/BS p=0.11).

To analyze the time-dependent frequency components of the response, we analyzed the recordings using a continuous Morlet wavelet transform. As previously, the transforms were averaged for all sweeps in every single experiment, and the frequency components prior to stimulus exposure were subtracted. The results are expressed as z-score (see Materials and Methods). **Figure 2C** shows a spectrogram of the response recorded at the OB after exposure to BS, where the maximum frequency of the oscillation suffers a shift towards lower frequencies as the response progresses in time.

Although in many recordings, the response extended past 5 s (sometimes reaching up to 10 s), the most robust and consistent responses were localized in the 1.5 to 4 s window. We calculated the average maximum frequency for that time span, shown in **Figure 2D**. All responses to AA, SE, and BS show the same frequency shift, starting at a higher frequency and then decreasing by about 2 Hz after 2 seconds. Interestingly, this analysis shows more evidently that BS evokes oscillations at a higher frequency than the other odorants.

### Spectral Decoding in OB, Vv and Dp

To confirm the notion that spectral power can separate odorant identity within olfactory areas (Schütt et al., 1999; Losacco et al., 2020), we performed a robust statistical approach based on spectral Multivariate Pattern Analysis (MVPA or “spectral decoding”;King and Dehaene, 2014). Spectral decoding allows us to obtain a measure of odorant discrimination without having to specify at which areas or frequency bands these differences emerge, while at the same time extracting subtle trial-by-trial neural differences that are undetected by standard averaging procedures (**Figure 2**)(Fahrenfort et al., 2018b). To do so, we trained a classifier to simultaneously distinguish the three types of odorants across the three regions (OB, Dp and Vv) (**Figure 3B**; see Materials and Methods). The above-chance classification accuracies imply that the relevant information about the decoded odorants is present in the oscillatory activity, implying spatially distributed olfactory processing and coding.

Interestingly, and contrary to the standard averaging statistical procedures described above, spectral decoding showed that information about the three odorant categories was reliably decoded above chance. In the multivariate case, where the three olfactory regions were combined (**Figure 3B**), a cluster-based permutation test showed a significant cluster of increased classification accuracy spanning several frequencies and time points (cluster p < 0.01; peak frequency: 6-10 Hz; time range: 1 to 9 secs). Similarly, in the univariate case, where decoding was performed on each area separately, we observed comparable results to the multivariate results (**Figure 3C**). These findings confirm that the spectral power of olfactory oscillations contains information about odorant identity in relevant olfactory areas, which can be revealed by spectral decoding.

### Nonlinear connectivity measures across the olfactory system

Functional connectivity – understood as coordinated interactions across brain areas – is a key element of several theories of perception across species(Fries, 2005; Vinck et al., 2023). While classic linear metrics of functional connectivity such as coherence investigate the temporal synchronicity between stereotypical patterns observed in the LFPs (i.e., neural oscillations), metrics based on information processing capture interactions that are not necessarily linear or oscillatory (King et al., 2013). Importantly, non-oscillatory or ‘aperiodic’ information is critical for establishing nonlinear connectivity when brain signals are highly complex(Vinck et al., 2023; Gelens et al., 2024).

### Information sharing between OB, Vv and Dp

Experimentally, nonlinear connectivity can be investigated by computing the shared information across brain regions, showing its robustness in discriminating perceptual representations (Canales-Johnson et al., 2020b, 2020c) and alertness states (King et al., 2013; Sitt et al., 2014; Imperatori et al., 2019). Based on these previous findings and on the spectral decoding results (**Figure 4**), we reasoned that a mechanism based on distributed information sharing across OB, Vv and Dp could be appropriate for distinguishing odor composition and intensity. Thus, we applied a method for quantifying nonlinear functional connectivity based on information theory: weighted symbolic mutual information (wSMI)(King et al., 2013). For each odorant category used here, we computed information sharing between regions in the frequency range of the oscillatory response (∼6-10 Hz; **Figure 4**). We first computed wSMI between OB and Vv (**Figure 4A**). Kruskal-Wallis test showed a significant interaction across odors (odors: TW, AA, BS, SE; X^2^ = 29.9, p<0.001). Post-hoc comparisons showed differences between odors and control (AA-TW: W = 5.85; p<0.001, SE-TW: W = 5.97, P<0.001, BS-TW: W = 4.80, p<0.001), and between two out of three odors (AA-SE: W = 3.52, p = 0.013; SE-BS: W = -3.78, p = 0.007; AA-BS: W = -1.81, p = 0.20). A similar interaction effect across odors was observed in wSMI between OB and Dp (**Figure 4B**; Kruskal-Wallis test: X^2^ = 28.0, p<0.001), with significant post-hoc effects between odors and control (AA-TW: W = 5.92; p<0.001, SE-TW: W = 5.49, P<0.001, BS-TW: W = 4.53, p<0.001), and between two odors (AA-SE: W = 3.46, p = 0.014; SE-BS: W = -3.78, p = 0.007; AA-BS: W = -2.29, p = 0.10). Interestingly, in the case of the wSMI between Vv and Dp (**Figure 4C**), we observed a significant interaction and simple effects across the three pairs of odors (Kruskal-Wallis test: X^2^ = 29.2 p<0.001; Post-hoc comparisons: AA-SE: W = 3.68, p = 0.009; SE-SB: W = -3.84, p = 0.007; AA-BS: W = -2.89, p = 0.032), and between odors and control (AA-TW: W = 5.38; p<0.001, SE-TW: W = 5.97, P<0.001, BS-TW: W = 4.90, p<0.001).

**Figure 4.**
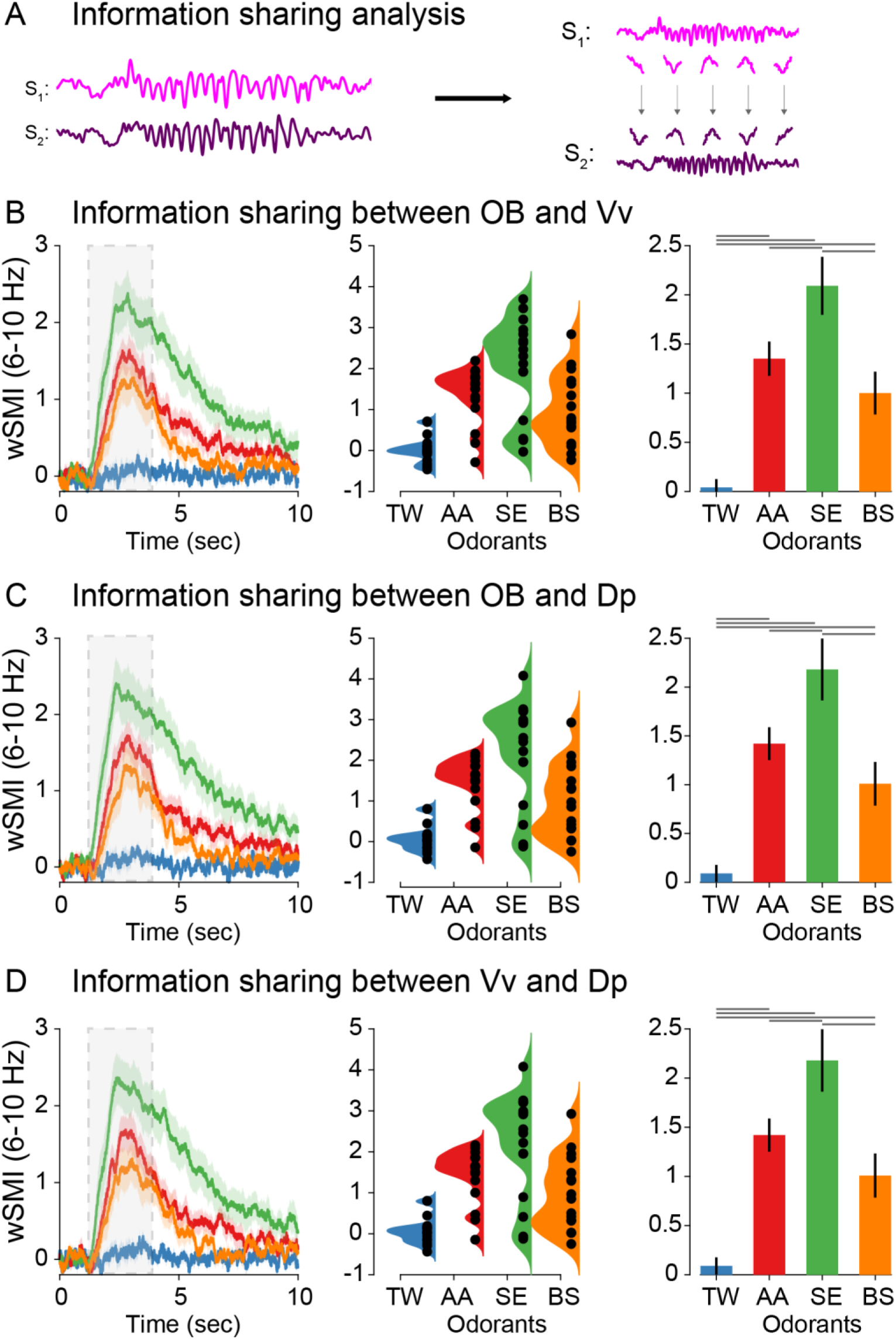
wSMI between OB-Vv, OB-Dp and Vv-Dp. **(A)** Schematic representation of wSMI analysis (i.e., weighted symbolic mutual information). Nonlinear transformations cause systematic relationships across different frequencies between signals, potentially in the absence of linear transformations. Thus, nonlinear functional connectivity measures quantify arbitrary mappings between temporal patterns in one signal (signal X) and another signal (signal Y). wSMI dynamics between OB-Vv **(B)**, OB-Dp **(C)**, and Vv-Dp **(D)** in the ∼6-10 Hz range for each odorant (left panel), the single-animal distribution of values during the 1-5 sec range (middle panel) and its group statistical analysis (right panel; *post-hoc* differences depicted as gray lines above the corresponding odorants). RANOVA revealed a significant interaction effect of wSMI between odorants and control (TW), and between most of the odorants after the *post-hoc* contrasts (see Results section).

### Information redundancy between OB, Vv and Dp

Robustness, understood as the ability of tolerating perturbations that might affect the system’s functionality, is a desirable characteristic for the olfactory system (Rabinovich et al., 2000), for instance to preserve odorant discriminability in the presence of noise, high background or highly variable odor plumes. Although wSMI provides relevant insight into the *amount* of information shared between brain regions to distinguish odorant identity, it does not provide any insight into whether the brain regions are processing the same or different information. Thus, in order to investigate robustness from an informational point of view, we need a more nuanced analysis capable of quantifying the *redundant* information that is being processed between regions (Ince et al., 2017). Observing redundancy between brain regions would suggest a common information processing pathways for odorant features (i.e., odor mix decoding).

Odorant information in two brain regions can be schematized in a Venn diagram (**Figure 5A**). The first quantity of interest is the overlap, termed *redundancy* (left panel, red area). Each inner circle represents the mutual information between pairs of odorant categories and their LFP signals (e.g. between AA and EPT) for an individual brain region (e.g. Vv). Conceptually, the term redundancy refers to the case in which the information conveyed by region A and region B is the same (e.g. Vv and Dp). If the variables are redundant, each brain region *alone* is sufficient to convey all the information about odorant category, and adding observation from the second brain region does not contribute additional information. On the other hand, the concept of *synergy* is related to whether region A and B convey *extra* information about odorant identity if both regions are considered jointly (right panel, blue area). Redundancy and synergy are reflected by *positive* and *negative* values of co-information (co-I), respectively (see Materials and Methods).

**Figure 5.**
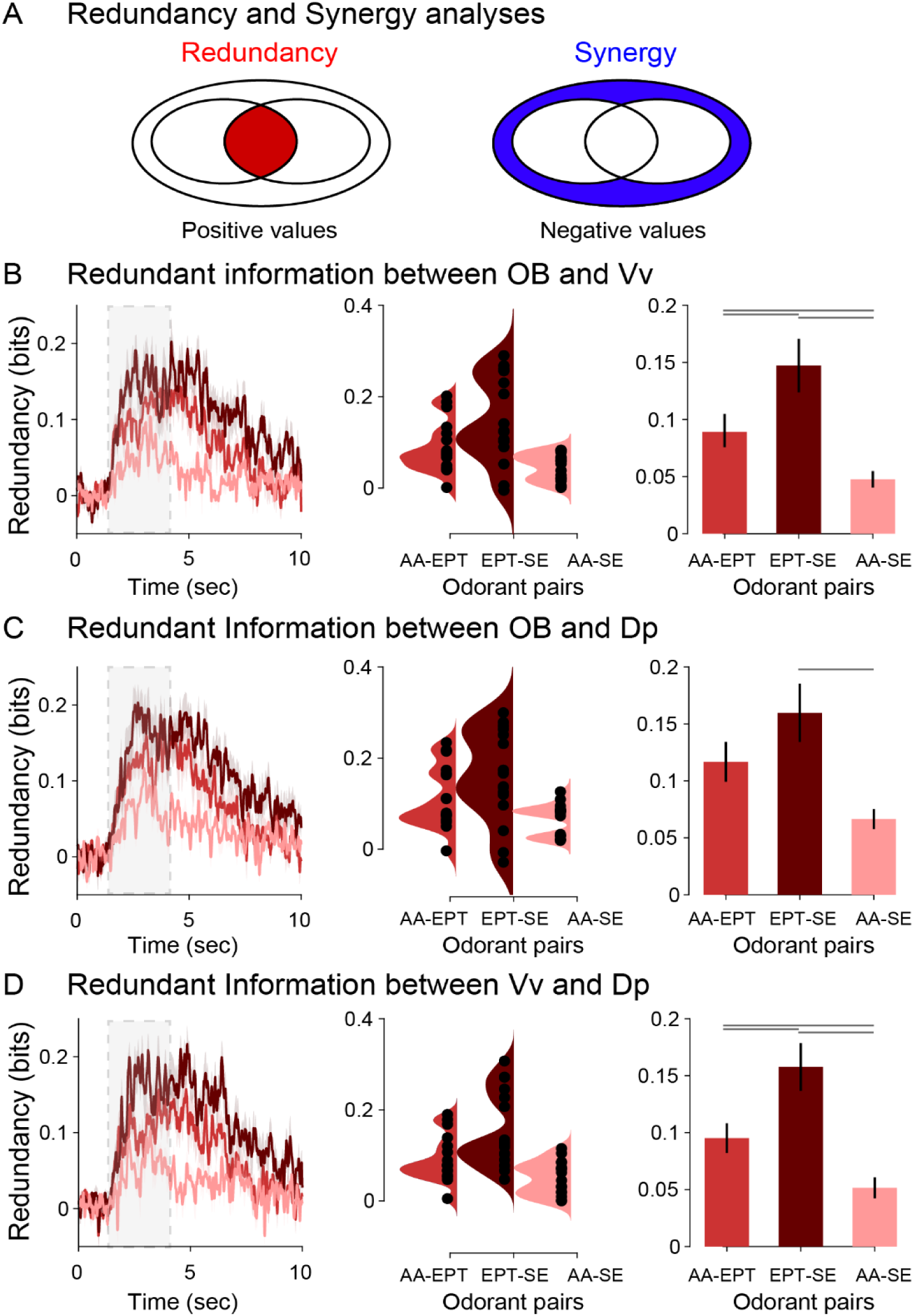
Information redundancy between OB-Vv, OB-Dp and Vv-Dp. **(A)** Schematic representation of redundancy and synergy analyses. Each inner circle represents the mutual information between LFP signals of a pair of odorants for an individual brain region, and the overlapping region represents the redundancy (red; left panel). The outer circle represents information that is synergistic (blue; right panel). Redundancy dynamics between OB-Vv **(B)**, OB-Dp **(C)**, and Vv-Dp **(D)** in the ∼6-10 Hz range for each odorant (left panel), the single-animal distribution of values during the 1-5 sec range (middle panel) and its group statistical analysis (right panel; *post-hoc* differences depicted as gray lines above the corresponding odorants).

Between pairs of brain regions, we computed the co-I about an odorant contrast (i.e. the difference between odorant A and odorant B) in the frequency range of the oscillatory response (∼6-10 Hz; **Figure 5**). We observed increased co-I in the post-stimuli period compared to the pre-stimuli period for all odorant pairs and between all brain regions, indicating redundant information processing (Ob-Vv: AA-EPT vs PSP, W = 13.86, p<0.001; EPT-SE vs PSP, W = 17.96; p<0.001; AA-SE vs PSP, W = 14.50, p<0.001; Ob-Vv: AA-EPT vs PSP, W = 12.78, p<0.001; EPT-SE vs PSP, W = 16.23; p<0.001; AA-SE vs PSP, W = 13.21, p<0.001; Ob-Vv: AA-EPT vs PSP, W = 17.01, p<0.001; EPT-SE vs PSP, W = 14.20; p<0.001; AA-SE vs PSP, W = 15.39, p<0.001).

Interestingly, the dynamics of redundancy was not the same for different pairs of odorants. Co-I was first computed between OB and Vv (**Figure 5A**). Kruskal-Wallis test showed a significant interaction (pairs: AA-EPT, EPT-SE, AA-SE; X^2^ = 13.2, p=0.001), and post-hoc comparisons showed differences between odor pairs (EPT-SE vs AA-EPT: W = 2.77, p<0.050; AA-SE vs EPT-SE: W = -4.74, p<0.001; AA-SE vs EPT-SE: W = -3.09, p = 0.029). A similar interaction effect across differences in odor pairs was observed between OB and Dp (**Figure 5B**; Kruskal-Wallis test: X^2^ = 9.81, p=0.007), with significant post-hoc effects in one contrast (EPT-SE vs AA-EPT: W = 2.13, P = 0.132; AA-SE vs EPT-SE: W = -4.37, p = 0.002; AA-SE vs EPT-SE: W = -2.35, p = 0.097). Interestingly, in the case of the redundancy between Vv and Dp (**Figure 5C**), we observed a significant interaction and simple effects across the three pairs of odor differences (Kruskal-Wallis test: X^2^ = 16.8, p <0.001; Post-hoc comparisons: EPT-SE vs AA-EPT: W = 3.09, p = 0.029; AA-SE vs EPT-SE: W = -5.49, p < 0.022; AA-SE vs EPT-SE: W = -3.25, p < 0.001).

Taken together, these results suggest that nonlinear interaction measures across olfactory areas reflect odorant representations. Moreover, this processing is odorant-specific, as the different odorants tested showed differences in the interaction between brain areas when assessed in pairwise fashion or in higher-order interactions.

### Linear connectivity measures across the olfactory system

In order to compare our results with canonical linear functional connectivity measures derived from spectral decomposition, we performed a phase synchronization analysis (**Figure 6**) using the weighted phase lag index (wPLI) due its robustness to volume conduction, common source and muscular artifacts (Vinck et al., 2011), and a coherence analysis (**Figure 7)**. In a similar manner as with wSMI analyses (**Figure 4**), we computed the phase synchronization between pairs of regions for each odorant in the frequency range (6-10 Hz) of the oscillatory response.

**Figure 6.**
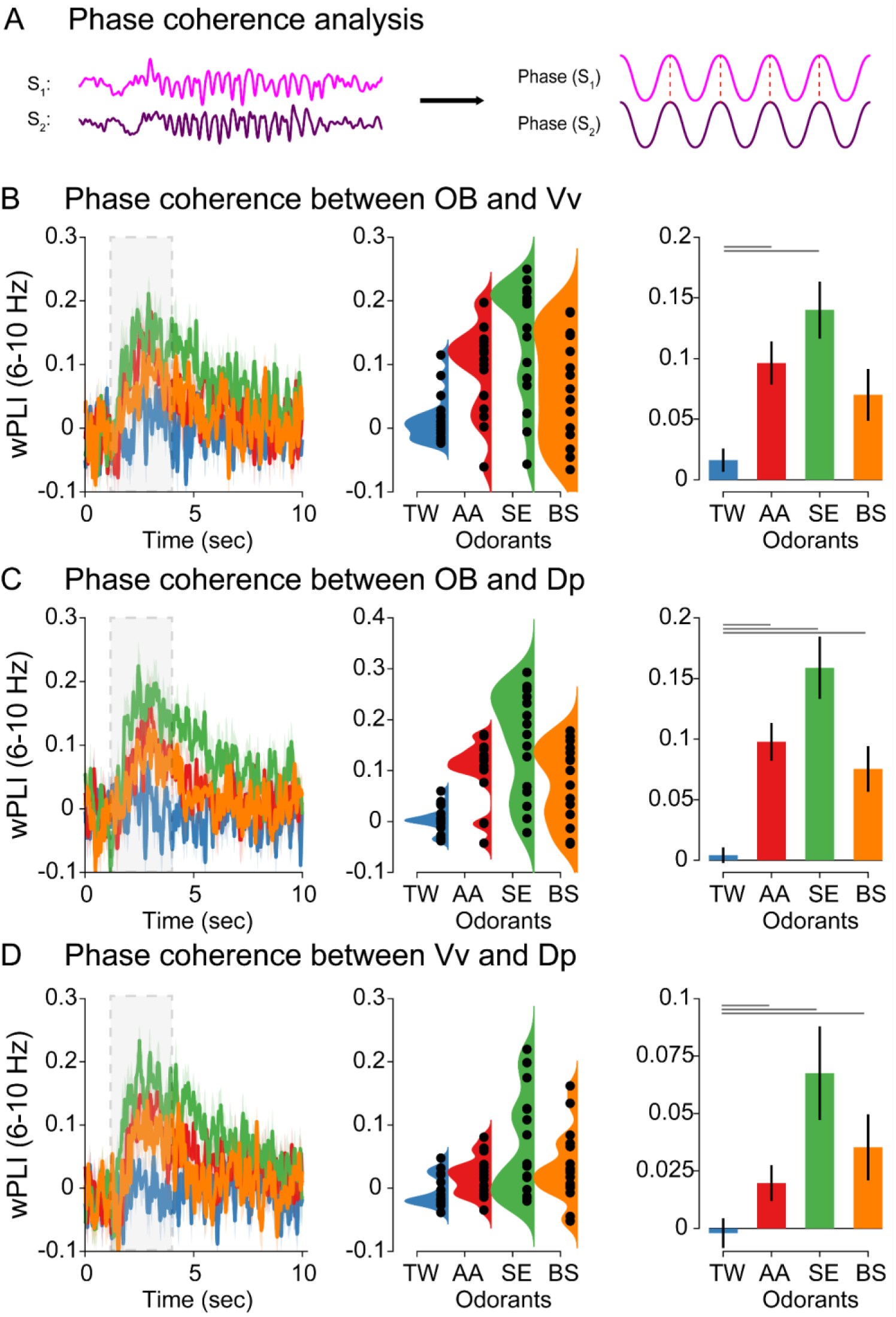
wPLI between OB-Vv, OB-Dp and Vv-Dp. **(A)** Schematic representation of wPLI analysis (i.e., phase coherence). After filtering the signal in a selected frequency range (see Materials and Methods) the instantaneous phase is computed for both signals S_1_ (pink) and S_2_ (purple), and the phase difference between them (red dashed line). In this example, phase difference between signals remains constant across time, representing a highly phase-coherent pair of neural signals. wPLI dynamics between OB-Vv **(B)**, OB-Dp **(C)**, and Vv-Dp **(D)** in the ~6-10 Hz range for each odorant (left panel), the single-animal distribution of values during the 1-5 sec range (middle panel) and its group statistical analysis (right panel; *post-hoc* differences depicted as gray lines above the corresponding odorants). RANOVA revealed a significant interaction effect in wPLI between odorants and control (TW), but no differences between pairs of odorants were observed after *post-hoc* comparisons (see Results section).

**Figure 7.**
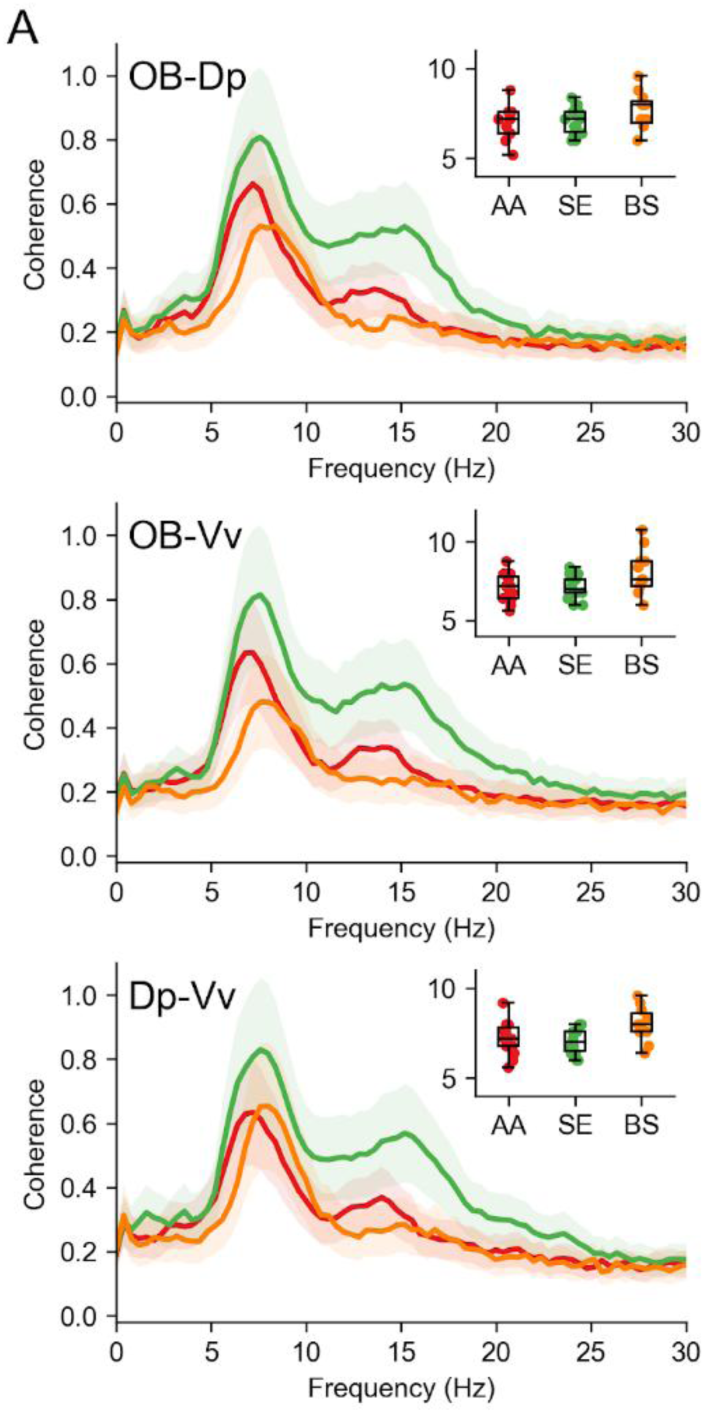
Coherence in the oscillatory response recorded in different regions. **(A)** Coherence spectra showing the mean (lines) and standard error (shades) of coherence detected at every frequency, in the pairs of regions indicated and after exposure to the odorants that evoked the most robust responses. For each experiment, a coherence spectrum was calculated and the experiments for each odorant were averaged. The inset shows the frequency where the peak of the coherence was detected.

### Phase synchrony across OB, Vv and Dp

In the case of wPLI between OB and Vv (**Figure 6A**), although the Kruskal-Wallis test showed a significant interaction across odors (X^2^ = 11.9, p = 0.008), post-hoc comparisons revealed no differences between odors (AA-SE: W = 2.13, p = 0.132; SE-SB: W = -2.18, p = 0.122; AA-SB: W = -0.42, p = 0.763), and only two odors were different from control (AA-TW: W = 3.67; p = 0.009, SE-TW: W = 4.21, P = 0.003, BS-TW: W = 2.61, p = 0.065). In the case of wPLI between OB and Dp (**Figure 6B**), a Kruskal-Wallis test showed a significant interaction across odors (X^2^ = 22.0 p<0.001) and between odors and control (AA-TW: W = -5.27; p < 0.001, SE-TW: W = -5.49, p <0.001, BS-TW: W = -4.05, p = 0.004). However, no differences were observed between odors (AA-SE: W = 2.07, p = 0.142; SE-BS: W = -2.13, p = 0.132; AA-BS: W = -0.80, p = 0.572). Similar results were obtained in the case of wPLI between Vv and Dp (**Figure 6C**; Kruskal-Wallis test: X^2^ = 22.3 p<0.001), with differences between odors and control (Post-hoc comparisons: AA-TW: W = 5.11; p <0.001, SE-TW: W = 5.54, p <0.003, BS-TW: W = -4.37, p = 0.002), but not between odors (Post-hoc comparisons: AA-SE: W = 2.45, p = 0.083; SE-BS: W = -2.66, p = 0.060; AA-SB: W = - 0.58, p = 0.679).

Altogether, these results suggest that the feature of the oscillatory response relevant for odorant discrimination is the shared information across regions and not the phase synchronization between them.

### Coherence across OB, Vv and Dp

Finally, we calculated the coherence cross-spectra between recording sites. **Figure 7A** shows the average coherence spectra of the responses recorded in presence of AA, SE and BS, for the three possible pairs between OB, Vv and Dp. All responses displayed a strong coherence with a peak around 7-8 Hz, the same frequency of the main oscillation. The median frequency of the maximum coherence (insets) was not significantly different between groups (Kruskal-Wallis test, OB-Dp H=2.62 p=0.27; OB-Vv H=4.4 p=0.11; Dp-Vv H=5.25 p=0.07). Interestingly, AA and SE also generated coherence in the 15Hz range, although this oscillation was not particularly visible in the frequency spectra. Most likely, this reflects harmonic components of the main oscillation frequency. Thus, the basic characteristics of odor-elicited oscillations allow for little or no discrimination between odorants. The only exception seems to be the higher frequency of oscillations elicited by bile salts.

## DISCUSSION

We have shown that odorant identity influences complex information-theoretic metrics, including information sharing and redundancy, across brain regions, highlighting nonlinear processing. In contrast, traditional linear functional connectivity measures, such as coherence and phase synchrony, exhibited minimal or no significant modulation by odorants. These results suggest that nonlinear interactions encoded by olfactory oscillations play a critical role in transmitting odor information within the teleost olfactory system, shedding light on the broader significance of nonlinear dynamics in sensory processing.

In the mammalian olfactory bulb, seminal studies conducted on behaving animals have demonstrated that neural activity at different spatiotemporal scales is modulated by animal behavior and experience apart from properties of the olfactory stimuli. For instance, in rabbits, amplitude-modulated patterns across 64 electrode arrays implanted in the OB exhibited highly context-dependent LFP oscillations during sniffing (Freeman and Schneider, 1982). Similarly, single-cell mitral and tufted cells were strongly influenced by contextual efferent inputs in behaving rats (Kay and Laurent, 1999). These studies suggest that LFP patterns in the OB contain a significant amount of non-primary sensory information, likely reflecting experience, the behavioral context, and associated information apart from primary characteristics of the stimulus.

If these studies account for the effects of context and behavior on olfactory processing, then what are the neural markers associated with odorant properties themselves? We sought to answer this question using anesthetized trout, allowing us to control confounding factors such as respiration and sniffing, movement, experience, and learning. Interestingly, spectral decoding and information-based functional connectivity analyses showed that the spatially distributed oscillations carry information relevant to odorant discrimination. We interpret the shared information conveyed in the LFP activity across OB, Dp and Vv as reflecting the complexity of the odor mixture. Arguably, stimuli formed by complex mixtures with diverse molecular structures should elicit a neural response containing a higher diversity of information patterns (i.e., higher signal entropy), resulting in an increase in information sharing between olfactory brain areas when they are co-activated.

The stimuli used in this study are blends of various odorants displaying different levels of complexity based on the quantity and diversity of their components (see Materials and Methods). Thus, the mixture of bile salts (BS) is composed of 4 compounds and the mixture of amino acids (AA) is composed of 5 amino acids. On the other hand, skin extract (SE), as a mixture resulting from the homogenization of a complete biological tissue, is composed of hundreds of compounds, many of them difficult to identify, conveying much greater complexity compared to AA and BS (Olivares, 2019). Although it cannot be directly concluded that the properties of the molecules that make up BS and AA determine greater or lesser complexity as both are mixtures of few compounds, the amount of their components can be arguably used as a proxy for structural complexity. Supporting this view, in all three pairs of regions, the wSMI results showed a correspondence between the quantity of components present in each odor, and the amount of information sharing between regions (**Figure 5B-D**). Thus, while SE showed the highest wSMI value (green bar; right panel), BS showed the lowest apart from control (yellow bar; right panel), and AA showed intermediate values (red bar; right panel). This correspondence between odor complexity – in terms of number of components – and the amount of information sharing between OB, Vv and Dp suggest that at least some of the information captured by wSMI encodes structural properties of the stimuli across the olfactory system.

To investigate nonlinear interactions between neural signals, we computed wSMI – an information-theoretic functional connectivity measure (King et al., 2013; Imperatori et al., 2019). Using realistically simulated EEG signals, we have recently demonstrated that wSMI can reliably detect highly nonlinear interaction across brain regions that phase synchronization (i.e., wPLI) was unable to detect (Imperatori et al., 2019). Why does the nonlinear connectivity measure (wSMI) capture odorant identity while the linear connectivity measure (wPLI) does not? The olfactory areas investigated in this study are thought to perform different roles during olfactory discrimination. While the OB is thought to decode the odorant’s molecular structure, higher olfactory centers such as Dp and Vv are thought to process more contextual and behaviorally relevant information such as feeding, reproduction, and danger-sensing (Yaksi et al., 2009). We propose that the associations between lower (OB) and higher (Dp and Vv) centers are established via nonlinear interactions across the olfactory system. Such nonlinear interactions are likely critical for pattern completion and separation (Vinck et al., 2023) and compute neural integration beyond the linear transmission of signals that dominated coherence and related measures like wPLI (Vinck et al., 2023; Schneider et al., 2021). It is worth mentioning that the current approach for measuring synchronization is based on bivariate measures (i.e., WPLI) and that potential synchronization distinguishing odorants might be encoded in multivariate functional connectivity patterns. Future studies employing multivariate synchrony (e.g., Safavi et al., 2023) could investigate this possibility.

Robustness is a functional consequence of degenerate systems, that is, systems conformed by structurally different elements capable of performing the same function (Edelman and Gally, 2001a; Whitacre and Bender, 2010; Marder, 2011a). Robustness is ubiquitous across many biological systems including neural circuits and networks (Tononi et al., 1999; Edelman and Gally, 2001b; Marder, 2011b). Crucially, robust systems are also capable of preserving their functions when exposed to changes in contextual circumstances, making them extremely resilient. From an evolutionary point of view, it is reasonable to conceive that selection processes such as those underlying the evolution of the olfactory system favor the development of robust systems. Phylogenetically, a mechanism preserving the implementation of the same function by different brain regions might serve a crucial evolutionary function: making olfactory discrimination quickly adaptable to changes in the environment. We have used information redundancy analyses to characterize the robustness of the teleost olfactory systems in implementing odorant discrimination. There is mounting evidence that redundancy in neural networks may provide various computational benefits, for example, enabling stable computations despite unstable neural dynamics (Druckmann and Chklovskii, 2012; Driscoll et al., 2017; Murray et al., 2017) and allowing the central nervous system to filter out unwanted noise (Moreno-Bote et al., 2014). Importantly, a recent study in the visual system has shown that oscillatory dynamics (i.e., narrowband gamma oscillations) convey primarily redundant information between visual areas, likely contributing to the robustness of visual representations (Roberts et al., 2025). Our finding of increased information redundancy across recording sites suggests that although neuroanatomically divergent, the underlying neural circuits of OB, Vv, and Dp can process the same information about odorant identity, supporting the idea that – at least under anesthetized conditions – the olfactory system of teleost is functionally robust.

General anesthesia dose-dependently suppresses neural signaling and can, therefore, be assumed to impact odor processing in different brain regions. Tricaine methanesulfonate (MS-222) is widely used as an anesthetic for aquatic organisms, particularly fish and amphibians, acting mainly through inhibition of voltage-gated Na+ channels, thus limiting nerve membrane excitability. The dosage of MS-222 used in the present study was comparatively low (10 mg/L) as it was applied together with an intramuscular injection of gallamine triethiodide (Flaxedil; 0.03 mg / 100 g body weight) as a muscle relaxant. For instance, a review on using MS-222 in fish indicates a dose of 60–100 mg/L in salmonids for major procedures (Carter et al., 2011). Therefore, its effects on odor detection and processing should have been limited. Several mammal studies have shown that general anesthesia does not significantly alter odor detection and processing. (Li et al., 2011)) reported that “levels of neural activities reached after the same odor stimulation had no significant difference.” Similarly, (Lang et al., 2013) and (Wachowiak et al., 2013), using intrinsic signals and GCAMP expression to detect odor representations in the olfactory bulb, found very similar response patterns between awake animals and different degrees of anesthesia. Furthermore, olfactory learning and memory processes in the olfactory bulb and forebrain continued under anesthesia (Nicol et al., 2014; Samuelsson et al., 2014). These studies suggest that odor processing is affected but not fundamentally altered by general anesthesia in the rodent olfactory bulb and forebrain, which could be extrapolated to salmonids. Together with the low level of anesthesia applied here, these antecedents support the general validity of our results for awake odor information processing.

In conclusion, our results reveal that odorant identity significantly modulates information sharing and redundancy across brain regions, emphasizing the role of nonlinear processing in olfaction. Conversely, traditional linear functional connectivity measures showed no significant modulation in response to odorants in the same brain regions and frequency ranges. These findings indicate that nonlinear interactions mediated by olfactory oscillations are essential for transmitting odor information in the teleost olfactory system, offering valuable insights into the broader importance of nonlinear dynamics in sensory processing.

## Contributions

Conceptualization: OS, ACJ. Data analysis: ACJ, PO, JO, VS. Visualization: ACJ, PO, JO. Trout recordings and surgeries: JO. Writing and editing: ACJ, OS, PO, JO. Supervision: OS, PO.

## Acknowledgments

We thank Dr. Martin Vinck for his helpful comments on the last version of this manuscript. OS is supported by ANID/FONDECYT Regular (1210790) research grant. PO is supported by ANID/FONDECYT Regular (1241469). ACJ is supported by an ANID/FONDECYT Regular (1240899) and ANID/FONDECYT Regular (1251273) research grants.

## Ethics

This study was approved by the bioethics committee of the Faculty of Science of Universidad de Valparaíso (CIBICA), certificate No. BEA 100-2017.

## Notes

### Competing Interest Statement

The authors have declared no competing interest.

### Summary of Updates

The abstract, introduction, and discussion were updated after peer review.

